# Can pseudotopological models for SMC-driven DNA loop extrusion explain the traversal of physical roadblocks bigger than the SMC ring size?

**DOI:** 10.1101/2022.08.02.502451

**Authors:** Biswajit Pradhan, Roman Barth, Eugene Kim, Iain F. Davidson, Jaco van der Torre, Jan-Michael Peters, Cees Dekker

## Abstract

DNA loop extrusion by structural-maintenance-of-chromosome (SMC) complexes has emerged as a primary organizing principle for chromosomes. The mechanism by which SMC motor proteins extrude DNA loops is still unresolved and much debated. The ring-like structure of SMC complexes prompted multiple models where the extruded DNA is topologically or pseudotopologically entrapped within the ring during loop extrusion. However, recent experiments showed the passage of roadblocks much bigger than the SMC ring size, suggesting a nontopological mechanism. Recently, attempts were made to reconcile the observed passage of large roadblocks with a pseudotopological mechanism. Here we examine the predictions of these pseudotopological models, and find that they are not consistent with some experimental data on SMC roadblock encounters. Particularly, these models predict the formation of two loops and that roadblocks will reside near the stem of the loop upon encounter – both in contrast to experimental observations. Overall, the experimental data reinforce the notion of a nontopological mechanism for extrusion of DNA.

## 1. Introduction

Higher order genome organization is largely dictated by structural maintenance of chromosome complexes (SMC) complexes^1–3^. SMC complexes including condensin, cohesin, and Smc5/6 in eukaryotes, share a ring-like architecture consisting of multiple subunits were two coiled-coil SMC arms and a intrinsically disordered kleisin constitute a large (∼35 nm) ring. Additional subunits attach to the kleisin and ATPase heads of the SMC subunits, rendering different functions to the different SMC complexes. SMC proteins organize the chromosome by DNA loop extrusion, as demonstrated by single-molecule in vitro experiments^4–7^. However, the mechanistic understanding of the coordinated movement of SMC subunits that underlie loop extrusion is still in its infancy.

The topology of DNA involved during loop extrusion has been a major point of discussion in recent years^1,8– 10^. Three possibilities have been proposed: i) topological binding, where the SMC ring opens to encircle the DNA, ii) pseudotopological loading, where a small nascent loop enters the SMC ring without opening, and where loop extrusion continues pseudotopologically by further extruding this inserted loop through the ring, iii) nontopological loading where the DNA does not enter the ring to extrude loop but instead binds at the outer interfaces of the SMC ring.

Recently, we showed that condensin and cohesin can bypass very large (even 200 nm) DNA-bound proteins and particles while extruding DNA loops^11^. Upon encounter of such a DNA-bound particle by the loop-extruding SMC complex, the particle transfers into the extruded DNA loop in the majority of cases. We even observed such passage of large particles by single-chain cohesins where all interfaces of the SMC subunits and the kleisin were covalently linked^5^. The passage of roadblocks bigger than the ring size of cohesin, even when the ring cannot open, clearly indicated that the extruded DNA does not enter the ring topologically or pseudotopologically. Instead, these experiments pointed to a nontopological mechanism for loop extrusion.

After these data appeared on bioRxiv in July 2021^11^, two new pseudotopological models were proposed by Shaltiel et al. (“hold-and-feed mechanism” ^12^) and by Nomidis et al. (“segment-capture model” ^13^). In the light of the roadblock experiments, the authors of these models attempted to explain the passage of the huge roadblocks within the context of their models.

Here we provide a detailed evaluation of these models with regards to the passage of large DNA-bound roadblocks upon encounter with a loop-extruding SMC complex. We spell out particular predictions that result from these models and evaluate whether or not these predictions are confirmed by experimental observations. We show that the predictions are not consistent with the experimental data for encounters of a loop-extruding SMC complex with DNA-bound roadblocks. The analysis thus does not support the pseudotopological models of loop extrusion, and instead point to a nontopological mechanism.

## 2. Description of the pseudotopological models

According to the models of Shaltiel et al.^12^ and Nomidis et al.^13^, SMC complexes entrap DNA pseudo-topologically, i.e., the two DNA strands that form the base of a loop pass through the ring structure formed by the SMC and kleisin subunits of these complexes. In view of the clear observation of the passage of roadblocks bigger than the SMC ring size into extruded loops in our experiments, the authors of these papers attempted to explain our observations within the context of their models. In other words, they provided an alternative interpretation of our data reported in ref.^11^. The authors of the papers argued that their model was not necessarily inconsistent with our observation that large roadblocks are able to enter extruded loops, even though these roadblocks are larger than the ring size of SMC complexes. If they are correct, it would question our conclusion that the data point to a nontopological mechanism for DNA loop extrusion by SMCs. Below, we evaluate this alternative explanation in the light of our experimental observations.

First, let us summarize the alternative model that Shaltiel et al. and Nomidis et al. proposed ^12,13^. We focus on the model of Shaltiel et al. who most explicitly pointed out the dynamics involved in the roadblock passage, according to their model, but the same considerations apply to the model by Nomidis et al. Without explicitly commenting on the detailed SMC conformational changes involved in these models (which also differ between these models), we describe how these authors attempt to explain the passage of roadblocks bigger than the ring size of SMCs. Figure 1 illustrates the model for roadblock passage by Shaltiel et al. Briefly, their model for loop extrusion involves the trapping of DNA in three chambers (I, IA, and II) in yeast condensin, where extrusion is proposed to proceed through a concerted action called hold-and-feed mechanism. DNA binding to Ycg1 and the associated kleisin region act as a “safety belt”. DNA loading into chambers I and II initiates extrusion of DNA. Two pseudo-topologically entrapped DNA loops are thus formed; one between chambers I and II, and the second one threaded between the SMC coiled coils. Upon ATP hydrolysis, these two pseudo-topological loops merge into one loop (loop-1), and the SMC starts extrusion of the next segment of DNA. Note that their figure illustrates only tiny loops, whereas in our experiments the encounter of a large roadblock with a DNA-extruding SMC is observed when the loop has been already significantly extended. Specifically, this means that the tiny black loop sketched in the bottom left of panel 1 of Figure S5 will in practice in our experiments be a very large loop of many kbp of DNA.

**Figure 1.**
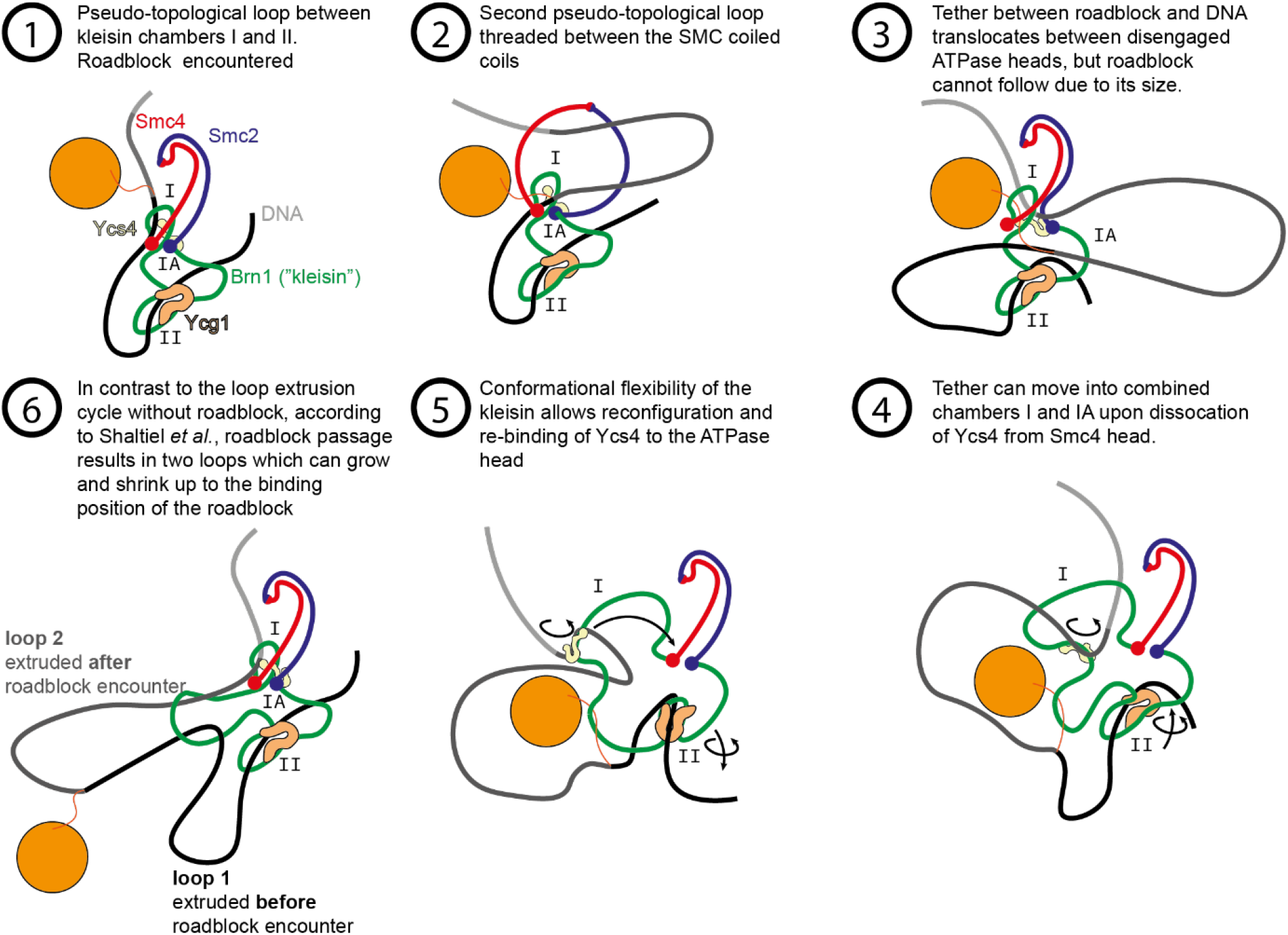
Model of the mechanism provided by Shaltiel *et al*. for roadblock passage into an extruded loop on the DNA. Figure adapted from Ref.12.

Rather than commenting on the plausibility of the various steps in these models, we here make predictions of what should be observed when a roadblock particle is encountered in the case that this alternative model would hold. Subsequently we turn to the experimental observations and evaluate whether or not these predictions are confirmed by the observations.

## 3. Predictions by the pseudotopological models

Specifically, the alternative models make the following predictions for the roadblock experiments:

### Prediction 1: Appearance of two loops in side-flow visualization

and

### Prediction 2: Roadblocks will reside near the stem of the loop upon encounter

When condensin encounters a large roadblock, the model predicts the existence of *two* loops (Figure 2A), not one, since the roadblock – with its size that cannot pass the ring structure – obstructs the merging of the two preformed loops described above. Next to the already formed extruded loop, a new second loop is therefore formed, where the latter loop has the roadblock particle attached.

**Figure 2.**
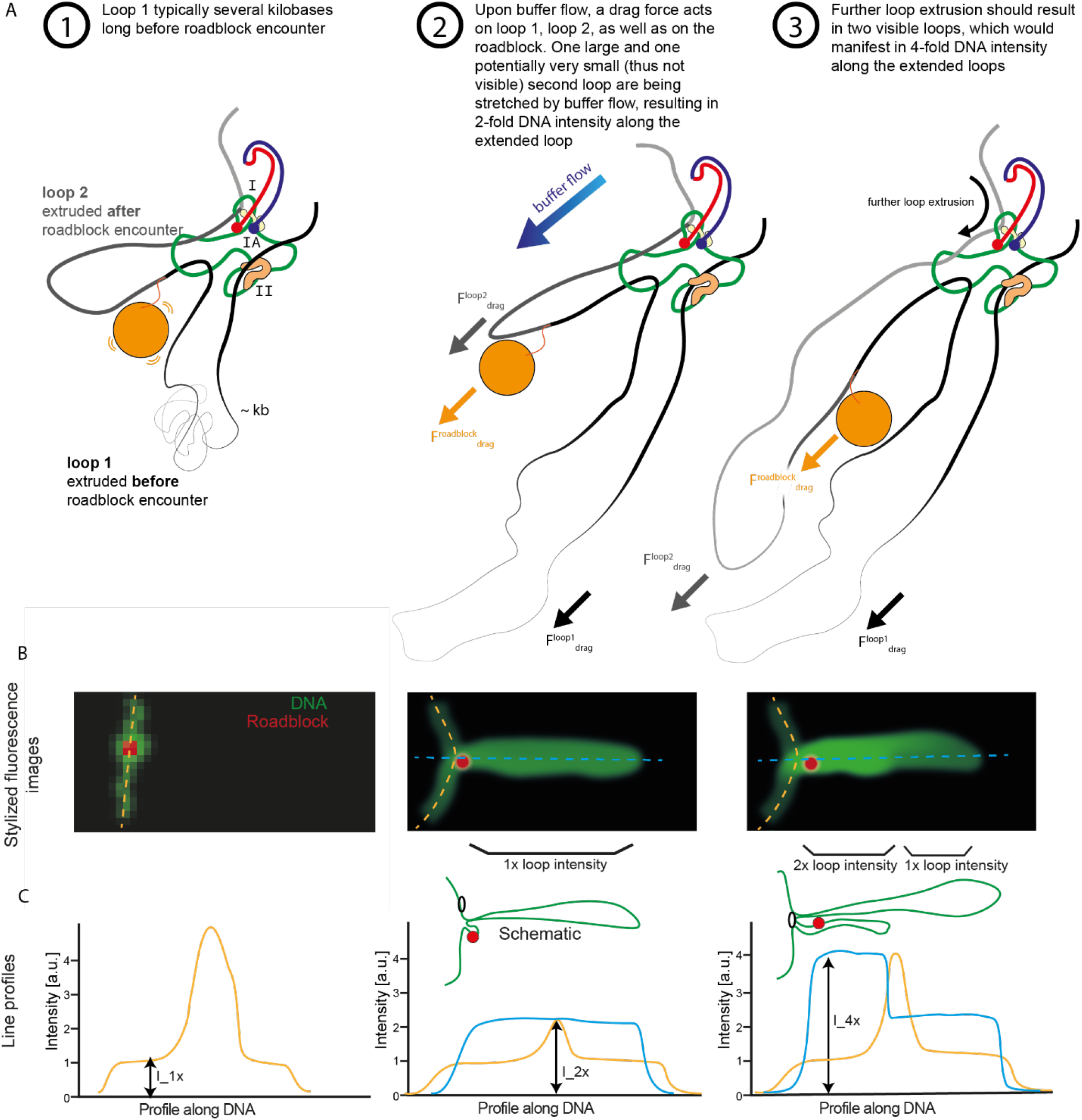
Two (not one) loops are associated with the model by Shaltiel *et al*. A. Presence of two loops during side-flow. B. Predicted stylized intensity images (not real experimental data) of the double tethered DNA (green) with the roadblock (red). C. Expected intensity profiles along the dashed lines in the images of panel B.

One could assume (as Shaltiel *et al* appear to do without further consideration) that the preformed loop-1 might slip into loop-2, to again form 1 joint loop. This however is not the case, as can be seen from considering the force balance: If side-flow is applied, several forces in action can be distinguished:

i. A drag force on the particle 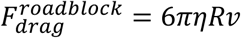, where the viscosity is *η* = 1 *mPa s* for a water-based buffer, *R* is the particle radius, and *v* the buffer flow speed. We experimentally measured the flow speed in side-flow experiments from occasionally observed beads that were not attached to DNA and were flowing through the field of view. We measured a flow speed of 79 ± 26 µm/s (n=3 independent experiments). The observed forces for different particle sizes are shown in Figure 3B, C and the values are below 0.15 pN for the largest 200 nm particles, and smaller for smaller roadblocks, down to 0.02 pN for a 30 nm particle.
ii. Drag force on loop-1 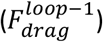. This can be estimated by calculating the tension on a flow-stretched DNA, as described by Pederson et al.^14^ Practically in our experiments, by the time that a roadblock encounters condensin, the size of loop-1 is observed to typically exceed 10 kilobases, which is associated with a drag force of value more than 0.27 pN (Figure 3C).
iii. Drag force on loop-2 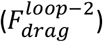. As loop-2 is a very small nascent loop (∼100 bp), the force acting on it is initially negligible, but it may become more significant as the loop continues to grow. The method to calculate the drag force acting on loop-2 is then analogous to loop-1.

It follows that upon encounter the drag force on loop-2 is nihil, while the drag force on loop-1 dominates and is larger than the drag force on the particle. As a result, the pre-encounter-formed loop-1 remains stable, while the particle remains localized close to condensin.

**Figure 3.**
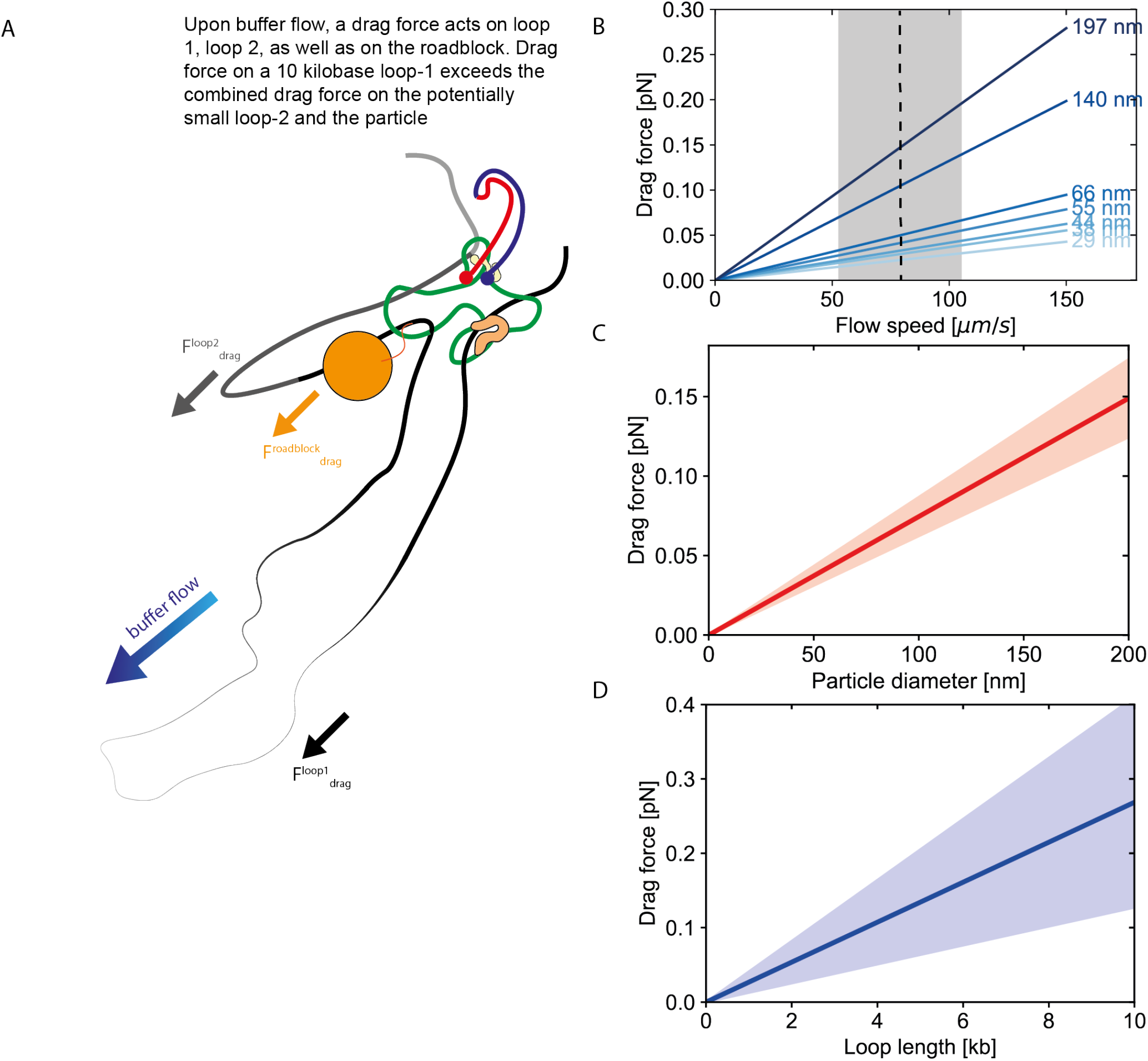
Drag forces on loops and particle associated with the model by Shaltiel *et al*. A) Schematic showing the blockage of the roadblock in presence of a large loop-1. (B) Estimated drag forces acting on particles of different sizes. The dashed line and grey area denote the experimentally measured flow speed of 79 ± 26 μm/s. C) The drag force on the roadblock (red, standard deviation based on uncertainty in buffer flow speed) versus particle size at the experimentally measured flow speed. D) Drag force on DNA loop versus loop length (blue line and shaded area denote mean and standard deviation, respectively. Standard deviations are based on the uncertainty in buffer flow speed (panel B) and drag coefficient^14^.

As loop extrusion proceeds, loop-2 grows and is slowly stretched alongside loop-1. Because of limitations in optical resolution, it is challenging to mutually distinguish the two loops spatially. However, the *intensity* of the two loops can be easily distinguished from that of a single loop by drawing intensity profiles across the loop during side-flow. Figure 2B and C illustrate what, according to the model of Shaltiel et al, would be expected in stylized intensity images and intensity profiles along the DNA for the pre-encounter and post-encounter loops. In summary, the model of Shaltiel et al predicts the observation of two loops that should be observed in a double intensity of that of the pre-encounter-formed loop-1 (prediction 1).

Furthermore, the model predicts that the roadblock, resides at the stem of the loop after encounter (prediction 2), while the second loop keeps growing. It is of interest to note two features connected to this prediction: First, the effects of spontaneous Brownian motion can be estimated to be small. *F* = −γ ν + δF(t) ≈ 1 *fN*; where γ = 6πηR, η is dynamic viscosity, R the radius of the particle, and the typical velocity of a 10 nm particle is ∼1 μm/s (http://book.bionumbers.org/what-are-the-time-scales-for-diffusion-in-cells/). The fluctuating force is on average zero, ⟨*δ*F(t)⟩ = 0, but the autocorrelation function can be given as ⟨*δ*F(t)*δ*F(t + *τ*)⟩ 2*γ*kT*δ*(*τ*), which leads to ⟨*δ*F(t)^2^⟩2*γ*kT ≈ (1fN)^2^. The Brownian forces to move the particle away from the SMC are thus very low, in the order of a few fN, so an appreciable spontaneous drifting away of the particle is very unlikely. Second, on long time scales, when the second loop would grow to a size comparable to the first loop (∼10 kbp), the drag force on the particle plus that on loop-2 may exceed the drag force on loop-1, which may initiate a slipping process. However, this would occur only rarely in our assay due to the finite total length of the DNA which limits the overall growth of the loop. Indeed, the typical loop size of loop at the moment of encounter (i.e. loop-1) was on average 12 kb in our experiments, while the added amount of DNA after that (i.e. loop-2 in the Shaltiel model) was 6 kb on average.

Summing up, according to the model, one would expect that the roadblock resides at the stem of the loop after encounter (prediction 2).

### Prediction 3: Loop size will increase while the MSD may stay constant after roadblock encounter

What happens when the loop extruding SMC encounters a roadblock on DNA? We note that the hold-and-feed model is not considering that loop extrusion will be blocked in this case, but rather that the roadblock ends up in a second loop. In the absence of side flow, upon encounter with the roadblock (as explained in the previous point), loop-2 with the roadblock particle grows through extrusion of DNA, but the particle stays close to the SMC due to the higher drag force on loop 1 than on the particle. This makes a specific prediction for the observation of simultaneously a continued growth of the loop size without, however, any significant increase in the MSD of the particle, as illustrated in Fig. 4A.

**Figure 4.**
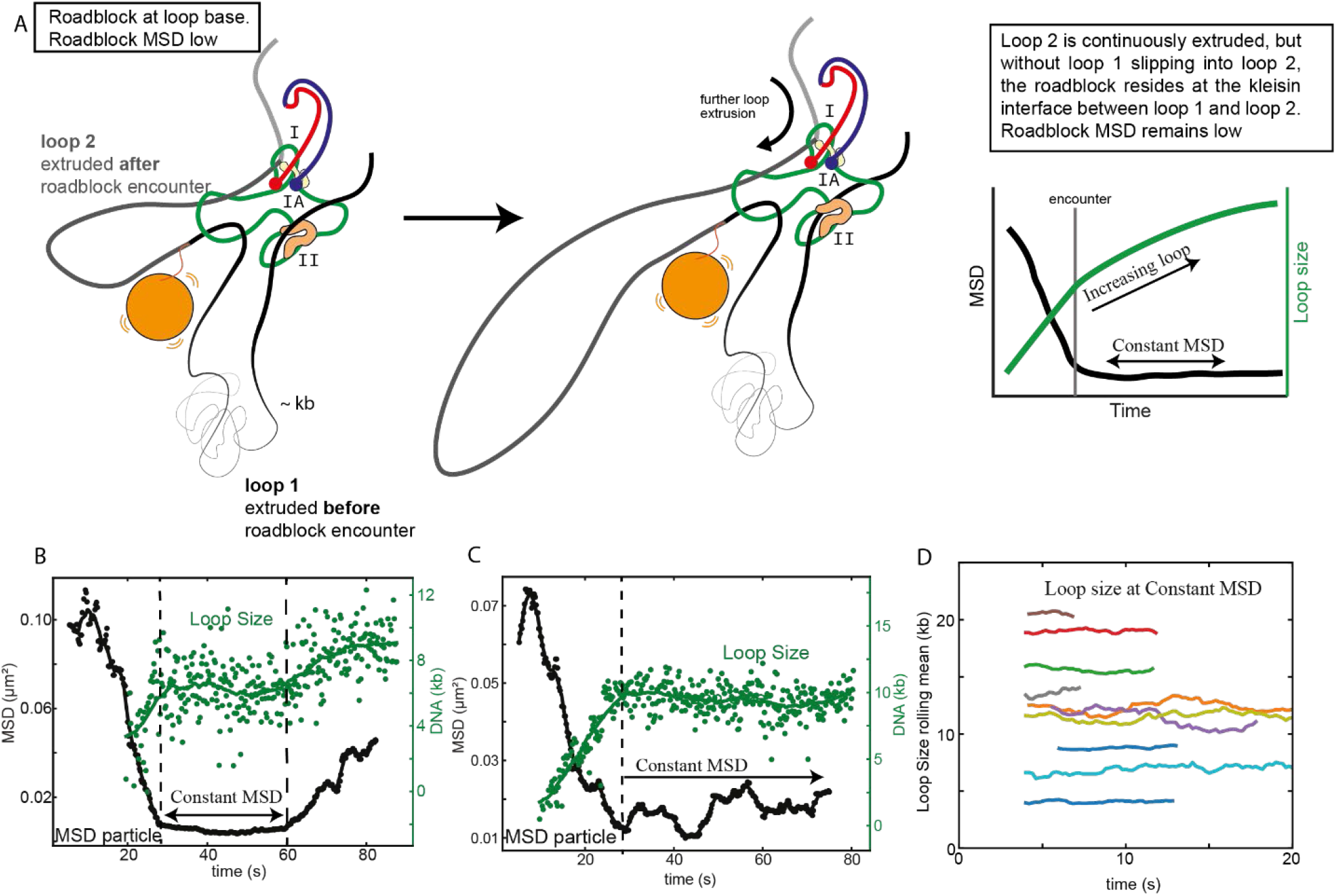
Test of prediction 3 associated with the model by Shaltiel *et al*. A. Correlation between DNA loop size and MSD of the particle after the encounter between condensin and roadblock, associated with the model by Shaltiel *et al*. B-C. Examples of kinetics of MSD (black) and loop sizes of extrusion (green) for events where the MSD remained low for a longer duration. D. Rolling mean of loop size with a window size of 20 points as a function of time in events where the MSD remained constant. Different colors represent different DNA molecules.

## 4. Experimental tests of the predictions made above

Our previous paper ^11^ presented an abundant set of experimental observations of encounters of a loop-extruding SMC complex and a DNA-bound roadblock. Here, we test whether or not the above predictions of the model by Shaltiel et al are confirmed by the experimental observations. For the latter, we refer to the published data^11^, but we here also present additional new data that were all acquired with the same methodology and under the same conditions as in Ref.^11^.

### Test of Prediction 1

To test whether or not the extruded DNA forms one or two loops when a roadblock is encountered, we measured intensity profiles of DNA at different stages of loop extrusion after a roadblock encounter, see Figure 5. The intensity of a single loop (Figure 5B) was deduced from the intensity profile on the DNA loop *before* the encounter between roadblock and the loop. The intensity of a single loop (i.e. I_2x) and the predicted putative double loop (I_4x) is shown in the lines in Figure 5C. We observe that the maximum intensity of the DNA equals the value of the intensity I_2x of a *single* loop while it never comes close to the predicted doubled loop intensity I_4x at any stage during loop extrusion at and after the encounter. From these side-flow visualizations, we thus did not obtain any indication of a doubled loop intensity (e.g. Figure 5A-D, F, G) – in contrast to the prediction of the alternative model by Shaltiel et al. Our data are therefore not supporting the key predictions made by this model.

**Figure 5.**
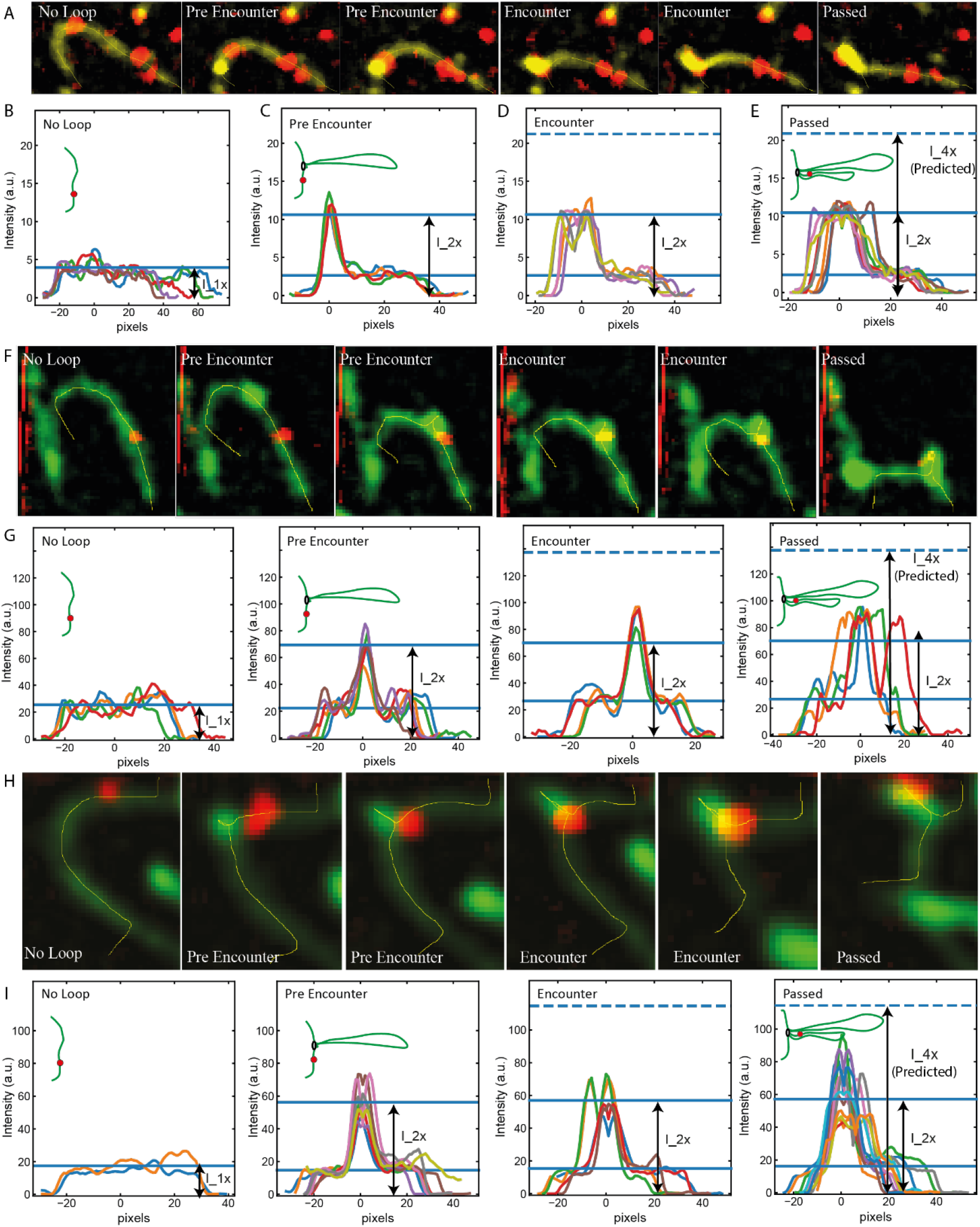
Test of prediction 1 associated with the model by Shaltiel *et al*. A) Snapshots of DNA and roadblock with sideflow representing different scenarios of intensity profiles in B-D namely No Loop, Pre Encounter, Encounter, and Passed the roadblock. Intensity profiles of DNA with a roadblock under sideflow before loop extrusion (B, No Loop), with loop but before encountering roadblock (C, Pre Encounter), at encounter when the roadblock is at the stem of the loop (D, Encounter), and after the encounter when the roadblock is inside the loop (E, Passed). Different colors represent different frames. F-I) Two more examples of the intensity profiles on DNA at different stages of the loop extrusion.

### Test of Prediction 2

The hold-and-feed model predicts that roadblocks will reside near the stem of the loop. This is, however, not observed. Instead, soon after the encounter, the roadblock is observed to be located within the extruded loop and separated from condensin or cohesin, see main figures from Pradhan et. al.^11^ and Figure 5A,F,H.

We can illustrate this in more detail by an example given in Figure 6. Here, the loop contained 24 kb of DNA before encounter of the roadblock. Upon encounter, the loop-extrusion speed did not abruptly change but loop extrusion continued monotonously at an approximately constant rate until the loop disrupted when it reached its final size of 30 kb. For this example, Shaltiel et al. predict the appearance of a second loop (loop-2) of 6 kb. However, no increased intensity is observed close to the stem of the loop, see Fig.6B-E. The model of Shaltiel et al. further predicts that the roadblock initially resides at the stem of the loop until the cumulative drag force on loop-2 and roadblock exceeds the drag force on loop-1. Note that the drag force on the 24 kb-long loop-1 is very large, ∼0.65 pN (Figure 3D), while the drag forces on the fully extruded loop-2 remain low at ∼0.16 pN and the drag force on the particle is estimated at ∼0.03 pN (Figure 3B-D). Hence, one does not expect any slippage of loop-1 into loop-2. The roadblock should therefore, according to Shaltiel et al., reside at the loop base while the loop continues to grow. However, the roadblock was experimentally observed to clearly travel continuously (and not suddenly as predicted for a potential slipping event) along one arm of the extruded loop (Figure 6A,F-H; and see main figures in the manuscript for more examples of condensin and cohesin). Quantitatively, after the encounter, the extruded loop enlarged from 1.6 µm to 2.1 µm under sideflow conditions (due to its increase from 24 kb to 30 kb). Each arm of the extruded loop in the final state thus contains 15 kb (half of the entire loop). For continuous loop extrusion into one loop, one would thus expect the particle to have moved to a distance of 6 kb/15 kb x 2.1 µm = 0.84 µm from the loop stem, which indeed is in excellent agreement with measurements of the ∼0.9 µm distance between roadblock and loop stem (Figure 6H). Yet another measure is to consider whether or not the particle moved into the loop with a speed comparable to the loop-extrusion speed. This would be expected in our scenario of continuous loop extrusion where the particle smoothly moves into the extruded loop, whereas Shaltiel et al. would predict the particle to stall at the loop stem and then suddenly speed up in a slippage event. Experimentally, we observed that the particle moved over a distance of ∼0.9 µm in 13 seconds, i.e., with a speed of ∼0.07 µm/s. This fits well with the loop-extrusion speed which for this example is ∼0.08 µm/s (since it is ∼0.5 kb/s (Figure S10I) and the 6 kb is extruded into 0.9 µm along one arm of the loop). We thus observe that the loop-extrusion speed is the same as the velocity with which the roadblock moves away from loop base.

**Figure 6:**
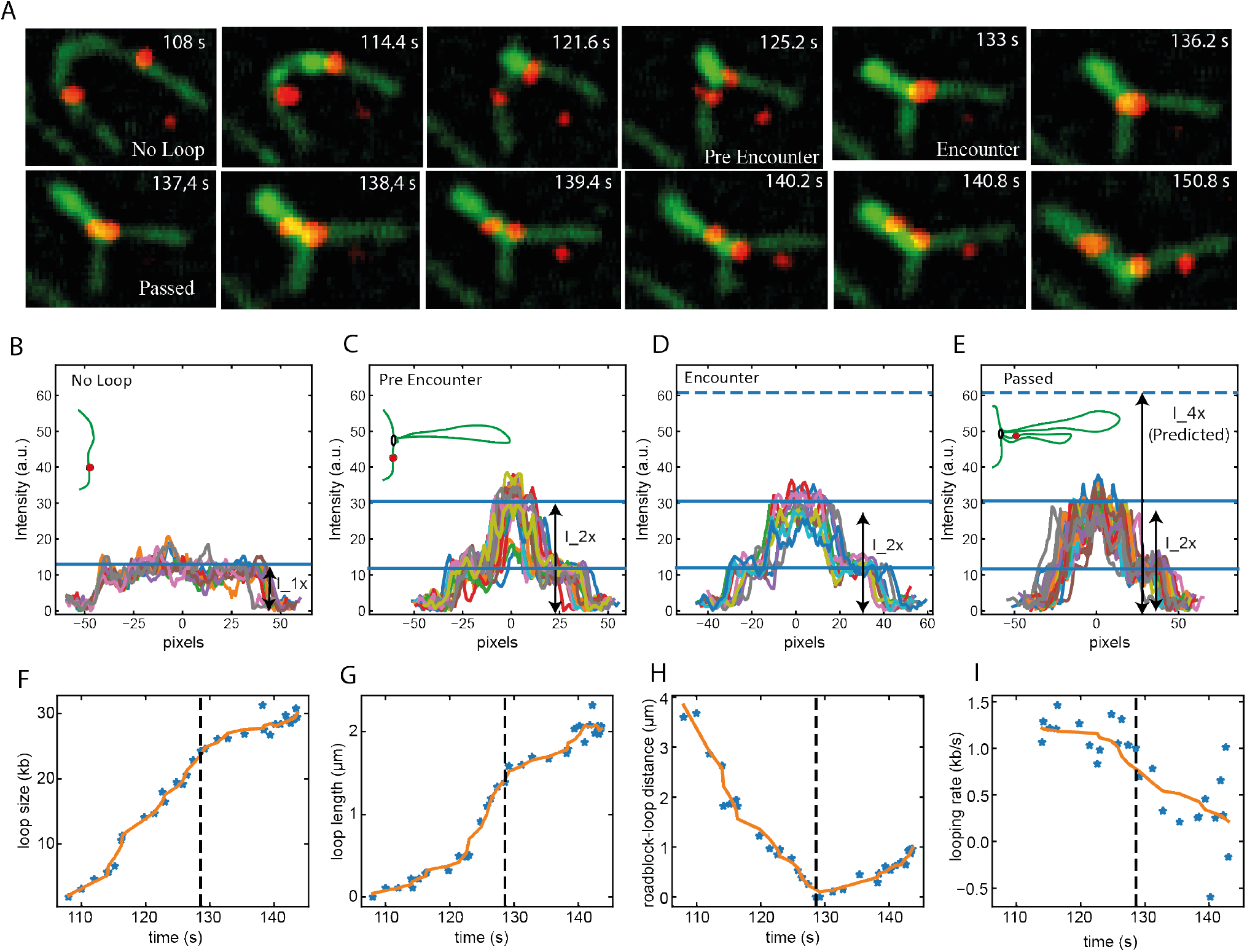
Test of prediction 2 associated with the model by Shaltiel *et al*. A) Snapshots of DNA (green) and a 39 nm-wide roadblock (red) with sideflow showing the continuous approach of the roadblock toward the loop stem before encounter (time points until 128 s) and the continuously increasing distance between roadblock and loop stem after encounter. B-E) Loop intensities during the time series. F) Loop size (kb) versus time. G) Loop length (μm) versus time. H) Distance (μm) between roadblock and loop stem versus time. I) Looping rate versus time. Dashed lines in F-I shows the time of encounter.

The data thus are in full accordance with continuous extrusion of the DNA and the particle into one loop, while the data are not in accordance with the suggested model by Shaltiel et al.

### Test of Prediction 3

How do the size of the extruded DNA loop and the MSD of the particle correlate? Throughout the manuscript (Figures 1-4 of Pradhan et al., 2021), we reported that we observed a strong correlation between an increase in loop size and an increase in MSD after the encounter with the roadblock. Here, we focus more in detail on stalling and passing events, where the MSD remained constant, and we searched for data that were compatible with prediction 3. In the model prediction, the loop size can grow while the MSD of the particle may stay constant. Note also that here we discuss experiments without an applied side flow. Figure 4B shows an example of what we called a stalling event where the MSD (black) remained constant for more than 10s. Clearly, the corresponding loop sizes during the period when the MSD was constant (e.g. between 30s and 60s in Figure S8B) was also found to be constant. This is the typical behavior. We examined all our stalling events (N=15), and we found that *none* of the events showed a concomitant increase in loop size (see Figure S8D for additional examples). The observation that the loop size *never* increased while the MSD remained constant is not consistent with one of the key predictions made by the model of Shaltiel et al.

## 5. Conclusion

We conclude that the predictions of the alternative pseudotopological models by Shaltiel et al. and Nomidis et al are not consistent with our experimental observations. Instead, our experimental data are fully consistent with a nontopological model of loop extrusion.

## Acknowledgments

We thank Benedikt Bauer, Theo van Laar, Wayne Yang, Je-Kyung Ryu, M. Tisma, A. Katan, I. Shaltiel, and C. Haering for discussions.

